# The Haemagglutinin Gene of Bovine Origin H5N1 Influenza Viruses Currently Retains an Avian Influenza Virus phenotype

**DOI:** 10.1101/2024.09.27.615407

**Authors:** Jiayun Yang, Mehnaz Qureshi, Reddy Kolli, Thomas P. Peacock, Jean-Remy Sadeyen, Toby Carter, Samuel Richardson, Rebecca Daines, Wendy S. Barclay, Ian Brown, Munir Iqbal

## Abstract

Clade 2.3.4.4b H5N1 high pathogenicity avian influenza virus (HPAIV) has caused a panzootic affecting all continents except Australia, expanding its host range to several mammalian species. In March 2024, H5N1 HPAIV was first detected in dairy cattle and goats in the United States. Since then, over 230 dairy farms across 14 states have tested positive, with zoonotic infections reported among dairy workers. This raises concerns about the virus undergoing evolutionary changes in cattle that could enhance its zoonotic potential. The Influenza glycoprotein haemagglutinin (HA) facilitates entry into host cells through receptor binding and pH-induced fusion with cellular membranes. Adaptive changes in HA modulate virus-host cell interactions. This study compared the HA genes of cattle and goat H5N1 viruses with the dominant avian-origin clade 2.3.4.4b H5N1 in the United Kingdom, focusing on receptor binding, pH fusion, and thermostability. All the tested H5N1 viruses showed binding exclusively to avian-like receptors, with a pH fusion of 5.9, outside the pH range associated with efficient human airborne transmissibility (pH 5.0 to 5.5). We further investigated the impact of emerging HA substitutions seen in the ongoing cattle outbreaks, but saw little phenotypic difference, with continued exclusive binding to avian-like receptor analogues and pHs of fusion above 5.8. This suggests that the HA genes from the cattle and goat outbreaks do not pose an enhanced threat compared to circulating avian viruses. However, given the rapid evolution of H5 viruses, continuous monitoring and updated risk assessments remain essential to understanding virus zoonotic and pandemic risks.

## Introduction

Influenza A virus (IAV) infection is a major global disease that continues to threaten wildlife, poultry production and human health. IAV is negative-sense, single stranded and segmented RNA virus. Wild aquatic birds act as the natural reservoirs of IAV, harbouring 16 HA and 11 NA subtypes [1]. The global poultry industry has been severely impacted by wild bird-driven spread of Avian influenza virus (AIV). Since the first detection of high pathogenicity avian influenza virus (HPAIV) H5 subtype in 1996, its precursor A/goose/Guangdong/1/96 (Gs/Gd/96) lineage has then rapidly evolved into more than 30 subclades [2, 3]. The current clade 2.3.4.4b H5N1 has become dominant since 2021 with noticeable genotypic change from predominant H5N8 to H5N1, which then spread globally[4]. Panzootic H5N1 viruses have then caused numerous outbreaks in avian species worldwide with unprecedented infections. In addition to birds, the clade 2.3.4.4b H5N1 viruses have also caused several outbreaks in mammalian species, such as in minks, foxes, domestic cats, and semi-aquatic mammals [5-8]. Human infection of HPAIV H5N1 remains sporadic. There have been 903 cases of human infections resulting in 464 deaths since 2003 [9], but only several dozen of these viruses have occurred in the recent panzootic.

However, with the increasing reports of clade 2.3.4.4b H5N1 infections in mammalian species, it is a concern that the virus could adapt to transmit efficiently by the airborne route which might facilitate infection of humans. While there were no reports of H5N1 virus infection in livestock like goats and cattle until March 2024, the Minnesota Board of Animal Health (MBAH) reported fatal infections of clade 2.3.4.4b H5N1 virus in newborn goats in February following the culling of HPAIV poultry housed on the same premises. The transmission may have been caused by shared husbandry and water source [10]. On 25^th^ March, 2024 United States Department of Agriculture (USDA) reported detection of clade 2.3.4.4b H5N1 virus in milk samples collected from symptomatic dairy cattle in Kansas and Texas [11]. A dairy farm worker with conjunctivitis was also found to be infected by a clade 2.3.4.4b H5N1 virus [12]. As of August 2024, the H5N1 virus has been further detected in cattle in thirteen states across the US. At present, the virus is thought to be predominantly transmitted through milking, and movement of infected animals or contaminated equipment between farms or states[13-15]. The virus transmission route to cattle has been proposed as mechanical resulting in mammary gland infection [16]. All these cases have been attributed to a single genotype B3.13 of clade 2.3.4.4b viruses, which so far has only been detected in the U.S [17]. In addition, based on Centers for Disease Control and Prevention (CDC) reports, there have been a further 13 human cases, all with mild symptoms but either attributed to contact with infected dairy cattle (n=4) or to poultry (n=9) that had acquired infection from dairy cattle and therefore are also “bovine” origin viruses. The outbreaks in livestock and humans highlight the urgent and ongoing need for risk assessment of the virus for potential spillover to humans.

The binding of viral HA to host cell surface sialic acid (SA) containing glycans is a prerequisite for AIV entry into host cells [18], as well as a determinant of host tropism. Influenza viruses from avian species generally have a preference towards α2,3-(SAα-2,3Gal)-linked SA, whereas human seasonal and pandemic influenza viruses have a preference towards α2,6-(SAα-2,6Gal) linked SA [19], this preference reflects the predominant SA linkages found in the avian gastrointestinal and human upper respiratory tract, respectively [20]. AIVs can switch receptor binding specificity to overcome host barriers and change host tropism by acquiring mutations in receptor binding site of HA [21]. Upon the attachment of HA to SA, AIV particles are internalised into endosomes and undergo conformational changes triggered by low pH between 5 to 6 in the maturing endosomes [22]. The acidity stability of HA plays a crucial role in virus transmission and host range. The HA of human transmissible viruses are generally more acid stable with a pH of ≤5.5 for membrane fusion, whereas many avian viruses have less stable HA proteins that fuse at pHs >5.5 [23]. This is thought to be related to the predominant transmission routes of these viruses, human seasonal viruses transmit by the airborne route and require more stability to survive the harsh microenvironments of aerosol particles and mildly acidic respiratory secretions. In addition to pH stability, thermostability of HA has also been shown to be important for AIV transmission in mammals, and may act as a proxy for pH stability[24].

The recent sustained outbreaks in dairy cattle farms raise credible concerns about the virus evolving to acquire mammalian adaptations and becoming zoonotic or potentially pandemic. HA protein is critical for receptor binding and its pH stability is crucial as an indicator of avian to mammalian host transmission. Therefore, we investigated the phenotype of the HAs of these viruses through receptor binding profiles, thermostability and pH stability comparing goat and dairy cattle H5N1 viruses compared with earlier H5N1 clade 2.3.4.4b viruses that circulated in Europe, as well as the emerging mutations in viruses associated with dairy cattle outbreaks. Our findings provide valuable insights into the ongoing evolution of H5N1 viruses in livestock and their potential veterinary and public health implications.

## Materials and Methods

### Ethics statement

All the procedures involving embryonated eggs were undertaken in strict accordance with the guidance and regulations of the UK Home Office under project licence number PP6471846. As part of this process, the work has undergone scrutiny and approval by the animal welfare ethical review board at The Pirbright Institute, incorporating the 3Rs and followed by ARRIVE (Animal Research: Reporting of *in vivo* experiment) guidelines for quality, reproducibility, and translatability of animal studies.

### Cells and viruses

Madin-Darby canine kidney (MDCK) cells, human embryonic kidney 293T (HEK-293T) cells, and African green monkey kidney epithelial (Vero) cells were acquired from (CSU) of The Pirbright Institute (TPI). The cells were maintained with Dulbecco’s Modified Eagle’s medium (DMEM) supplemented with 10% fetal calf serum (FCS) (Gibco) with 5% CO2 at 37 °C. The HA and NA sequences were acquired from GISAID (Table S1) and synthesised by GeneScript and cloned into a bidirectional pHW2000 vector. The polybasic cleavage site of H5 HA PLRERRRKR/GLF was replaced with monobasic PLGTR/GLF cleavage site for the study to be conducted at Containment Level 2 (CL2) laboratories. The internal segments of the RG viruses were from laboratory adapted strain A/Puerto Rico/8/1934 (H1N1) (PR8). The viruses were rescued by previously described protocol [35]. The rescued viruses were propagated in 9 to 10-day old embryonated hen’s eggs.

### Site-directed mutagenesis (SDM)

SDM plasmids were generated using single-site mutagenesis QuikChange II (Agilent) kit following manufacturer’s instructions. The primers (5’ to 3’) were synthesised at Merck.

T143A forward: gctcacccctagtgatgcttcatgatttggccagg,

T143 reverse: cctggccaaatcatgaagcatcactaggggtgagc;

Q154L forward: gcttgtccatacctgggagcaccctcc,

Q154L reverse:ggagggtgctcccaggtatggacaagc;

D171N forward: atctttattgttgggtatgcattgttctttttgataagccacacc,

D171N reverse: ggtgtggcttatcaaaaagaacaatgcatacccaacaataaagat;

A172T forward: aatgtggtgtggcttatcaaaaagaacgatacgtacccaacaataaagata,

A172T reverse: tatctttattgttgggtacgtatcgttctttttgataagccacaccacatt;

P174Q forward: gcttatcaaaaagaacgatgcataccaaacaataaagataagctacaataata,

P174Q reverse: tattattgtagcttatctttattgtttggtatgcatcgttctttttgataagc;

I178V forward: gattagtattattgtagcttacctttattgttgggtatgcatcgttcttt,

I178V reverse: aaagaacgatgcatacccaacaataaaggtaagctacaataatactaatc;

Q234R forward: cttccacgttgcccgtttactctggatctagtagctatttttgg,

Q234R reverse: ccaaaaatagctactagatccagagtaaacgggcaacgtggaag.

### Bio-layer interferometry (BLI)

Propagated viruses were purified through 30% and 60% sucrose gradient cushion at 27,000 rpm for 2h at 4□. The purified viruses were normalised by enzyme-linked immunosorbent assay (ELISA) to a known standard of 100 pmol, quantifying viral nucleoprotein (NP) expression [36]. The standardised 100pmol virus was then incubated with 10 μM oseltamivir carboxylate (Roche) and 10 μM zanamivir (GSK) against avian-like sugar analogue 3SLN and human-like sugar analogue 6SLN, and receptor binding affinity was tested by Octet® R8 system (Sartorius) using streptavidin biosensors (Sartorius). Virus binding affinity was normalised to fractional saturation and the concentrations of sugar loadings [37].

### Antisera preparation

Chicken polyclonal antisera against Sct477/21 and Eng598/22 were prepared as previously described [38]. In brief, the viruses were propagated in embryonated hen’s eggs and then inactivated with 0.1% (v/v) β-propiolactone (BPL). The inactivated viruses were then passaged three times in embryonated hen’s eggs and checked by haemagglutination assay. The inactivated virus was then concentrated by ultracentrifuge at 27,000 rpm for 2h. Three-day-old specific pathogen free (SPF) chickens were inoculated with 1024 haemagglutinating unit (HAU) of concentrated inactivated virus mixed with oil emersion adjuvant (Montanide; Seppic) at a adjuvant : virus 7:3 ratio. A boost dosage was given after 10 days. The inoculated chickens were bled at 18, 25 and 38 days post-inoculation for HI validation.

### Haemagglutinin inhibition (HI) assay

Haemagglutinin inhibition assay were performed following World Health Organisation (WHO) guidelines [39]. Haemagglutinin inhibition (HI) test was carried out by incubating two-fold serially diluted chicken antisera with 4HAU for 1h, and then added 1% chicken red blood cells (RBCs). The HI titres were recorded after 30min incubation at room temperature.

### Virus neutralisation test (VNT)

Chicken antisera were heat-inactivated at 56°C for 30min and then serially diluted with DMEM. The diluted antisera were incubated with viruses at a concentration of 100 TCID50/mL in a 37°C incubator for 1h. Monolayered MDCK cells were washed with PBS and infected with the virus-antisera mixture for 1h at 37°C. After three washes with PBS, the cells were maintained in DMEM supplemented with N-tosyl-L-phenylalanine chloromethyl ketone (TPCK) trypsin for three days. Neutralisation titres were determined by staining the cells with crystal violet.

### Syncytium formation assays

Syncytium formation assays were performed as previously described [40]. In brief, viruses were titrated in Vero cells by using anti-nucleoprotein (anti-NP) mouse monoclonal antibody and horseradish peroxidase-labelled rabbit anti-mouse immunoglobulins (Dako). Monolayered Vero cells were then incubated with 1.0 multiplicity of infection (MOI) of each virus for 1h at 37°C, the cells were then washed with PBS and incubated at 37°C for 15h. The Vero cells were then treated with 3.0μg/mL TPCK trypsin in DMEM for 15min and followed by incubation with PBS ranging from pH 5.2 to 6.0 at 0.1 pH increment for 5min. The cells were then maintained with DMEM+10%FCS for 3h. The cells were then fixed with acetone: methanol (1:1 ratio) and stained with Giemsa stain (Sigma-Aldrich). Stained cell images were taken by EVOS XL imaging system at 400μm (Life Technologies).

### Thermostability assay

Thermostability assay was tested as previously described [40]. Viruses were diluted with allantoic fluid and normalised to 32HAU/50μL. The viruses were then incubated in Thermal cycler (Bio-Rad) at 50°C, 50.7°C, 51.9°C, 53.8°C, 56.1°C, 58.0°C, 59.2°C, and 60°C and 4°C as control for 30min, the HA titres were then determined by Haemagglutination assay.

### Haemagglutinin structure prediction

The crystal structure of H5 HA was acquired from Protein Data Bank (https://www.rcsb.org/) with assession number 4JUL. HA structure modelling and prediction were visualised using SWISS-MODEL and visualised and annotated in PyMol version 4.6.

### Statistical analysis

Data were analysed and visualised by GraphPad Prism 10.0 (GraphPad Software, USA). Significance was determined by two-way ANOVA multiple comparisons and Ordinary one-way ANOVA Dunnett’s multiple comparisons test. Levels of significance (p) are denoted as: 0.01<p<0.05 having one asterisk, and 0.001<p<0.01, 0.0001<p<0.001 and p<0.0001 having two, three or four asterisks, respectively. p>0.05 was considered not significant.

## Results

### HA genes from cattle and goat H5N1 viruses retain high binding affinity to avian receptor

The HA genes from genotype B3.13 associated with early Texas cattle outbreak and Minnesota goat outbreak, A/dairy cattle/Texas/24-008749-001-original/2024 and A/goat/Minnesota/24-007234-003-original/2024 (referred to as TX-Cattle and MN-Goat, respectively) were selected for comparison with the HA genes of the dominant H5N1 genotypes AIV09 and AIV48 from the UK [25, 26], A/chicken/Scotland/054477/2021 and A/chicken/England/085598/2022 (referred to as Sct477/21 and Eng598/22, respectively). For each HA, mutagenesis was performed to remove the multibasic cleavage site, and recombinant viruses were rescued by reverse genetics with 6 internal genes from PR8 and homologous NA. Receptor binding profiles against simple sialylated avian receptor analogue α-2,3-sialyllactosamine (3SLN) and human receptor analogue α2,6-sialyllactosamine (6SLN) were assessed by bio-layer interferometry (BLI). Both TX-Cattle and MN-Goat only showed binding affinity to 3SLN without binding to 6SLN (Figure 1A and 1B). Consistent with our previous study, HA genes of Sct477/21 and Eng085/22 also showed binding to 3SLN only (Figure 1C and 1D) [27]. A pandemic 2009 H1N1 virus, A/England/195/2009 (referred to as Eng195/09), and a zoonotic H7N9 virus, A/Anhui/1/2013 (H7N9) (referred to as AH1/13) were used as assay controls, with Eng195/09 binding only to 6SLN, and AH1/13 binding to both 3SLN and 6SLN (Figure 1E and 1F) [28]. The relative binding affinities of the panel viruses to 3SLN were also compared, with Sct477, Eng085/22 and TX-cattle showing comparable binding to 3SLN, while MN-Goat showed 17-fold stronger binding to 3SLN than Sct477/21 (Figure 1G).

**Figure 1.**
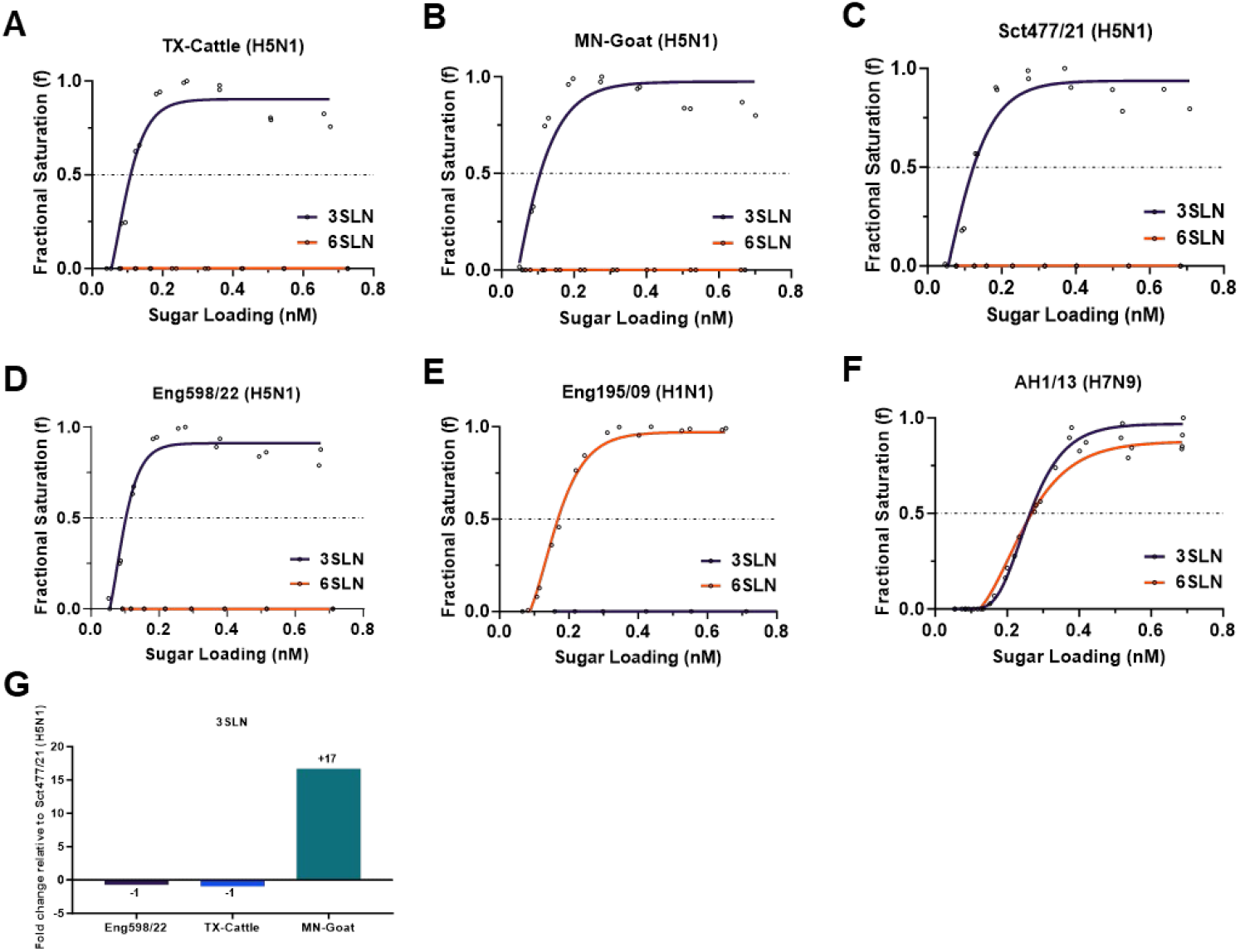
Receptor-binding of the panel clade 2.3.4.4b H5N1 viruses. The receptor binding of the H5N1 viruses was determined by BLI using avian-like receptor analogue 3SLN and human-like receptor analogue 6SLN. (A) TX-Cattle (H5N1), (B) MN-Goat (H5N1), (C) Sct477/21 (H5N1) and (D) Eng598/22 (H5N1), (E) Eng195/09 (H1N1) as assay control, which shows binding to 6SLN only. (F) AH1/13 (H7N9) as assay control, which shows binding to both 3SLN and 6SLN. The fold change of 3SLN relative to Sct477/22 (H5N1) is indicated in (G).

### HA genes from cattle and goat H5N1 viruses showed membrane fusion at pH 5.9, and goat H5N1 virus showed higher thermostability

We then tested the pH that triggered membrane fusion of the virus panel using a syncytium formation assay. Vero cells were infected with the viruses and treated with PBS at different pH values ranging from 5.3 to 6.0. pH of membrane fusion was assigned based on the highest pH at which syncytia formation was observed [29]. Both TX-Cattle and MN-Goat showed syncytium formation at pH 5.9 and below as indicated by red arrows but not at pH 6.0 and Sct477/21 and Eng598/22 at pH 5.8, but not at pH 5.9 (Figure S1). Thus both UK H5N1 viruses were assigned a membrane fusion pH of 5.8, while TX-Cattle and MN-Goat HA fused slightly more readily at pH 5.9 (Figure 2A).

**Figure 2.**
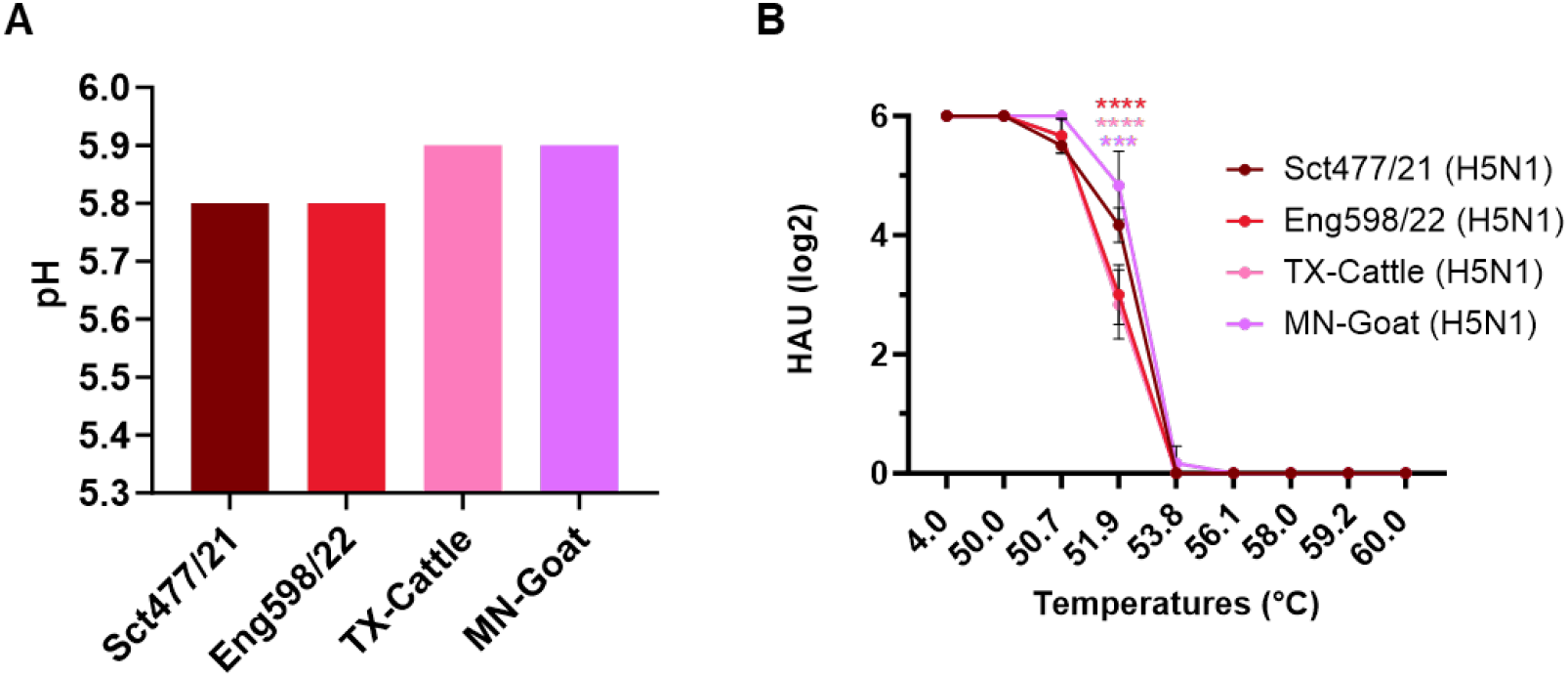
The HA stability of the panel clade 2.3.4.4b H5N1 viruses. (A) pH fusion of the panel H5N1 viruses. Syncytium formation assays were used to evaluate the fusion pH of panel viruses by infecting monolayered Vero cells in a range of pH treatments. (B) Thermostability of the panel H5N1 viruses. Virus loadings were standardised to 64 HAU and then incubated at the indicated temperatures for 30min. Haemagglutination assay was then performed after the incubation. The experiment was performed independently for three times, and levels of significance are shown in the figure, 0.0001<P<0.001 was given three asterisks and P<0.0001 was given four asterisks.

The thermostability of the virus panel was assessed by haemagglutinin assay. The viruses were standardised to 64 hemagglutinin units (HAU) and then heat-treated in a thermocycler from 50°C to 60°C for 30 minutes or kept at 4°C as a control to test their thermostability. The HAU of the virus panel remained unchanged at 50°C but were completely abrogated at 56.1°C. MN-Goat virus showed the highest thermostability, retaining about 4 times higher HAU at 51.9 C, than TX-Cattle and Eng598/22 (Figure 2B).

### HA genes from cattle and goat H5N1 viruses retained similar antigenicity compared to UK 2.3.4.4b viruses

The HA protein is immunodominant and mutations therein modulate antigenicity. We then examined the antigenic differences between the panel of viruses using antisera raised in chickens against Sct477/21 and Eng598/22. TX-Cattle and MN-Goat viruses showed ∽3log2 (8 fold?) reduction in HI titres in comparison to homologous titres against Sct477/21 and Eng598/22 (Figure 3A). The antigenicity of the virus panel was also assessed by virus neutralisation test (VNT) in MDCK cells. There was no significant change in virus neutralisation titre (Figure 3B) among the panel viruses (Figure 3B). These data together suggest that the panel of viruses were antigenically similar.

**Figure 3.**
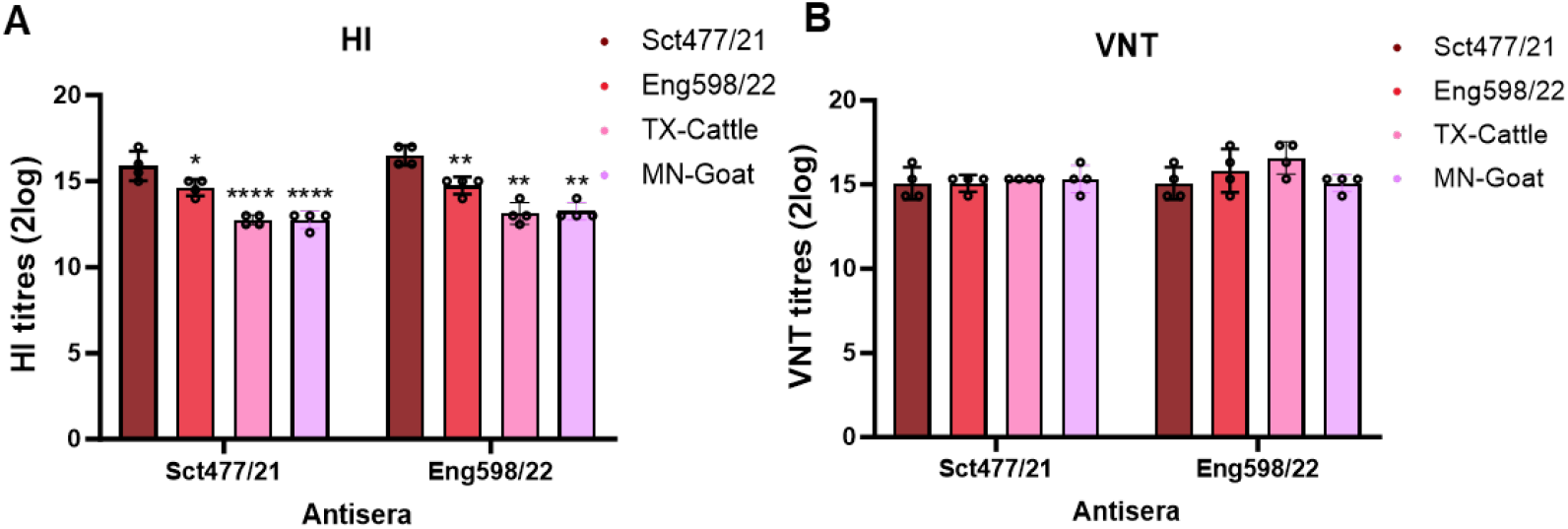
Antigenicity assessment between the panel H5N1 viruses. (A) HI titres of the panel viruses using Sct477/21 and Eng598/22 antisera. HI assays were performed using indicated antisera in four repeats. Levels of significance are shown in figures, 0.01<P<0.05 was given one asterisk, 0.001<P<0.01and P<0.0001 were given two, and four asterisks, respectively. P>0.05 was considered not significant. (B) Virus neutralisation of the panel H5N1 virus. Neutralisation tests were performed using the VNT assay with indicated antisera in monolayered MDCK cells for 3 days.

### The emerging HA mutants from cattle H5N1 HPAIV retained binding affinity to avian receptors

Based on 490 available virus sequences uploaded to Sequence Read Archive (SRA), we identified seven emerging mutations in bovine viruses mapping to the HA head domain that could potentially influence receptor binding. These mutations were T143A, Q154L, D171N, A172T, P174Q, I178V and Q234R by immature H5 numbering (T127A, Q138L, D155N, A156T, P158Q, I162V and Q218R by mature H5 numbering; T132A, Q142L, D159N, A160T, P162Q, I166V and Q222R by H3 numbering) with frequencies of 9.1%, 0.4%, 0.6%, 1.8%, 0.8%, 0.6% and 0.2%, respectively (Figure 4A). We engineered each of these mutations into recombinant virus using reverse genetics and tested their sialic acid specificity. We found that all the mutant viruses exhibited detectable binding to 3SLN only, but not 6SLN (Figure 4B-4H), suggesting the most prevalent mutations in the cattle viruses are not currently resulting in a switch towards human receptor binding. We further compared the fold change in 3SLN binding between mutant viruses to wild-type (WT) TX-Cattle, T143A, D171N, I178V and Q234R showed increased binding to 3SLN by 6-, 28-, 4- and 84-fold, respectively. Mutants with Q154, A172T and P174Q had decreased binding to 3SLN by 3-, 199- and 5-fold, respectively (Figure 4I).

**Figure 4.**
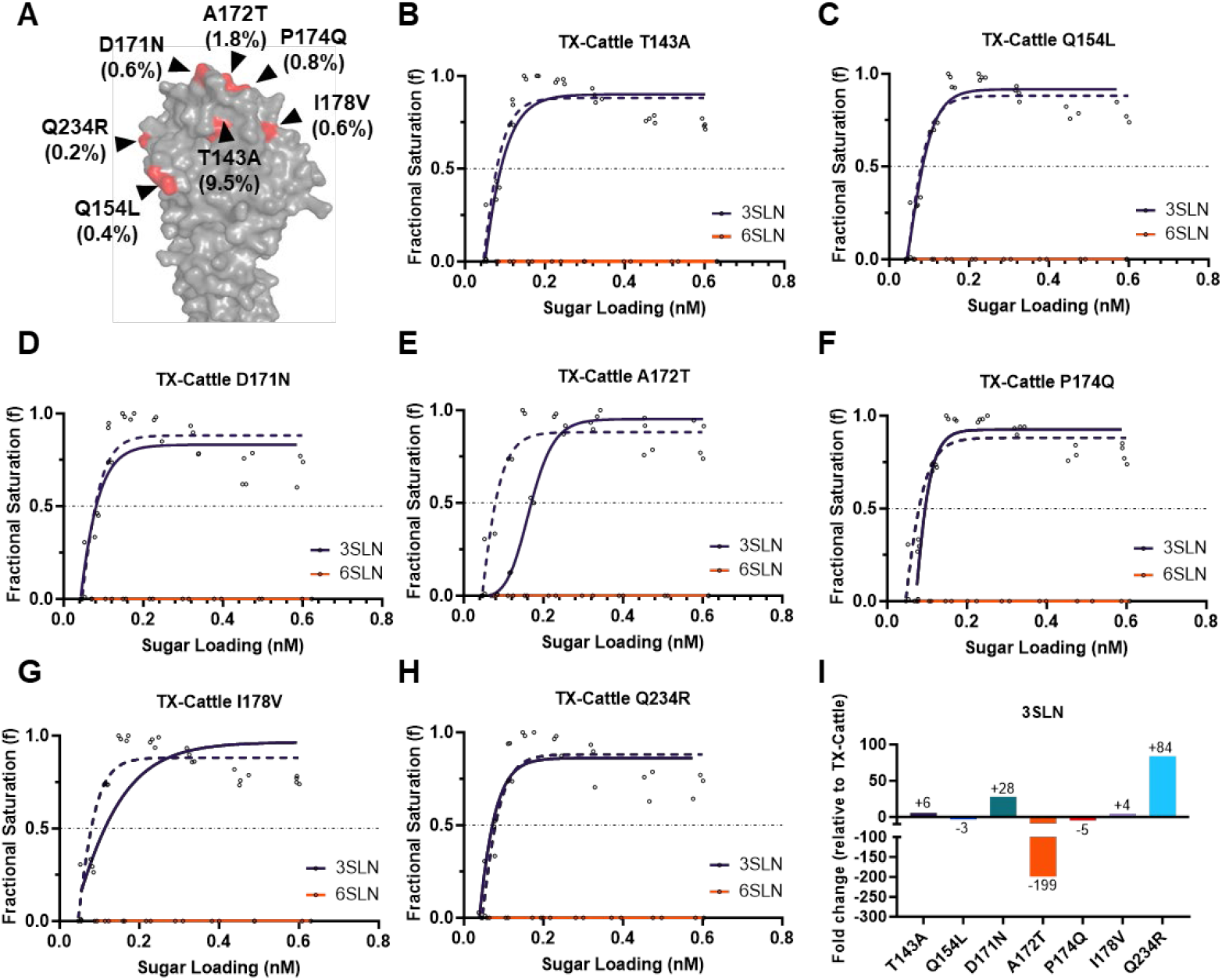
Receptor binding profiles of the emerging mutants from cattle outbreaks. (A) Seven emerging mutations (highlighted in red) are in the HA1 region of HA. Their frequencies (in brackets as percentage) in cattle outbreaks are shown below the mutations. BLI was used to assess the receptor binding to avian-like receptor analogue 3SLN and human-like receptor analogue on the following mutant viruses (in solid lines) compared to wild type TX-Cattle (in dotted line): (B) TX-Cattle T143A, (C) TX-Cattle Q154L, (D) TX-Cattle D171N, (E) TX-Cattle A172T, (F) TX-Cattle P174Q, (G) TX-Cattle I178V and (H) TX-Cattle Q234R. The relative fold change of binding affinity to 3SLN relative to the wild type TX-Cattle is shown in (I) with indicated fold change.

### The emerging HA mutants retained a fusion pH above 5.8, and do not confer any change in thermostability

We tested the pH threshold of membrane fusion for the cattle mutant viruses. Apart from Q154L which did not fuse until pH dropped to pH 5.8, the fusion pH of all six other mutant viruses, T143A, D171N, A172T, P174Q, I178V and Q234R, remained unchanged at pH 5.9 compared to the WT TX-Cattle (Figure 5A). The thermostability of the mutant viruses was also tested following heat treatment. A172T, P174Q, and I178V showed lower HAU compared to WT at 50.7°C and 51.9°C (Figure 5B), suggesting lower thermostability. All the other four mutations, T143A, Q154L, D171N and Q234R showed comparable stability to WT (Figure 5B).

**Figure 5.**
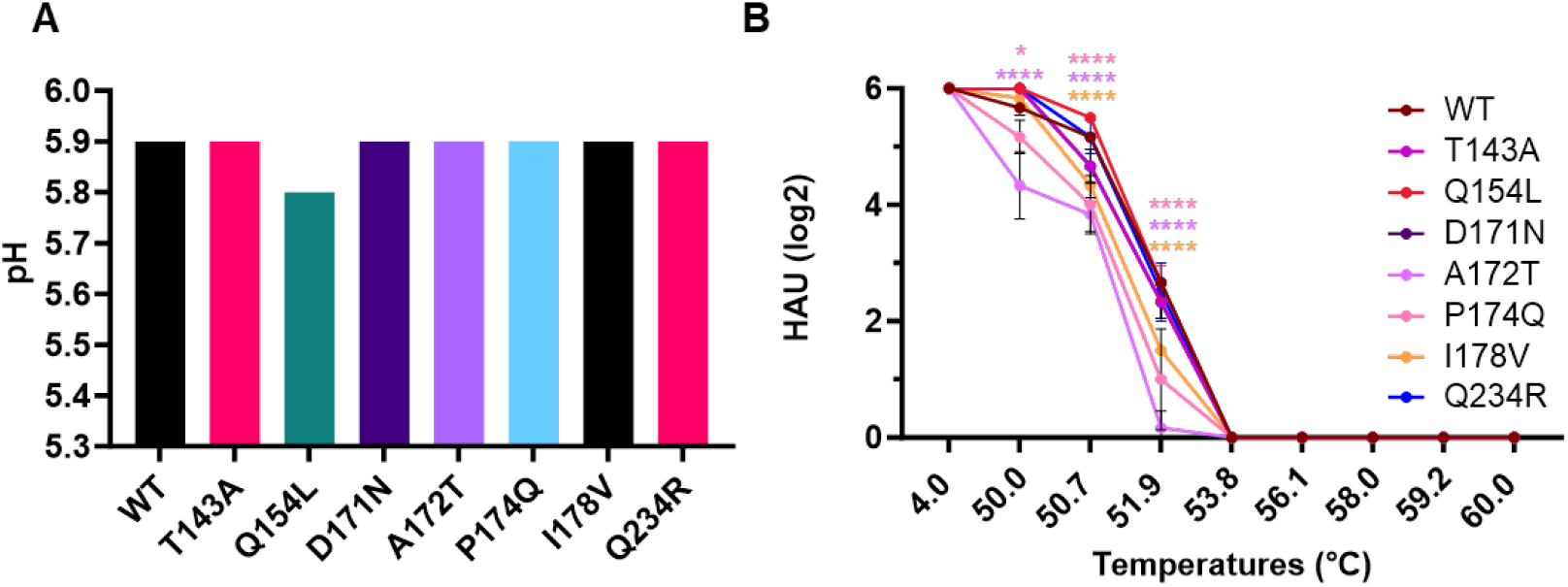
HA stability of the emerging mutants from cattle outbreaks. (A) Required pH for membrane fusion of the mutant viruses. Syncytium formation assays were used to evaluate the fusion pH of mutant viruses by infecting monolayered Vero cells in a range of pH treatments. (B) Thermostability of the H5N1 mutant viruses. Sixty-four HAU of the mutant viruses were heat-treated at the indicated temperatures for 30min. Haemagglutination assay was then performed after the heat-treatment. The experiment was performed independently for three times, and levels of significance are shown in the figure, 0.01<P<0.05 was given one asterisk and P<0.0001 was given four asterisks.

## Discussion

This study investigated HA properties known to be important in conferring avian to mammalian transmissibility of influenza A viruses including receptor binding, pH of fusion and thermostability in an effort to understand the propensity of the clade 2.3.4.4b virus to cross into cattle and the ongoing risk of zoonotic or pandemic infection in humans. Our BLI data showed the H5 HA of cattle H5N1 HPAIV from Texas virus had receptor binding to avian-receptor analogue 3SLN only. The finding aligns with several studies examining the receptor binding profiles of the human isolate from Texas dairy farm worker and cattle isolate from Ohio, which consistently showed virus binding to avian-type SA receptors only [30-32]. Only one recent study showed that the New Mexico cattle H5N1 virus displayed dual receptor binding to both human and avian receptors by solid-phase ELISA assay [33]. The only amino acid difference between the HA sequences of cattle Texas H5N1 virus and the New Mexico H5N1 virus was at S336N (S320N by H3 numbering or S323N by H5 numbering). This mutation seems unlikely to contribute to changes in receptor binding suggesting the discrepancy could be due to methodological differences. In addition, we also tested the receptor binding of the virus from the Minnesota goat outbreak and the virus also did not show receptor binding to human-receptor analogue 6SLN. Overall, these results indicate that the HA genes from the recent cattle and goat outbreak viruses retained binding to avian receptors.

The pH stability and thermostability of the viral HA protein are important for host range and virus transmission. Avian influenza viruses generally have a higher pH of membrane fusion (pH>5.5) compared to human seasonal influenza viruses, which require a lower pH of membrane fusion (pH≤5.5) to transmit efficiently by the respiratory airborne route [28]. In this study, we tested the fusion of cattle and goat H5N1 viruses, our data demonstrated syncytium formation at a pH of 5.9, indicating the virus currently has limited potential for human-to-human transmission, an essential pre-requisite to raise the level of pandemic risk. This may also suggest that this virus is currently not spreading efficiently by airborne route between cattle. To support this, although ferrets directly inoculated with the cattle H5N1 virus exhibited noticeable clinical signs and the virus replicated efficiently in the respiratory tract, eyes, and brain. no aerosol transmission was observed from infected ferrets to naïve ferrets [33], a surrogate for human respiratory droplet transmission.

It has been shown that both avian α-2,3-SA and human α-2,6-SA linkages are distributed in the mammary gland of cattle, suggesting cattle could potentially be a novel ‘mixing vessel’ for AIV or be an host in which adaptive mutations towards human receptor binding might be selected [34]. To further test HA adaptations that could potentially increase or switch receptor binding to human-like receptor analogues, we examined the receptor binding profiles of 7 emerging mutations, T143A, Q154L, D171N, A172T, P174Q, I178V and Q234R, that map close to, or within the receptor binding domain (RBD) of HA. Our data showed that H5 HA mutants bearing any of the 7 mutations still exclusively bound the avian-like receptor analogue. We further tested whether there was any change in the required pH for membrane fusion of the mutant viruses. Q154L showed only a slight change of membrane fusion at pH 5.8, while the other 6 mutations retained WT pH of fusion. Altogether, these findings indicate that despite the presence of α-2,6-SA linkages in cattle, no HA mutations in the cattle H5N1 viruses have yet acquired receptor binding affinity to the human receptor, and the mutant viruses are unlikely to have enhanced human-to-human airborne transmissibility due to retaining a high pH of fusion of 5.8-5.9.

In summary, we conducted a rapid risk profile of the receptor binding properties and fusion pH of ‘original’ dairy cattle, emerging HA mutants in dairy cattle, and goat H5N1 viruses. None of the tested viruses exhibited binding to the human receptor analogue 6SLN and they all retained a high membrane fusion pH ≥5.8, suggesting the currently circulating bovine H5N1 viruses are unlikely to be able to efficient transmit between humans. However, it remains to be determined whether further HA mutations will emerge if the virus continues to infect and spread amongst dairy cattle, with continuing opportunities to adapt to bovines. Addressing this question will contribute to a better understanding of the H5N1 virus and its adaptation in cattle. Continued surveillance and risk assessment of circulating clade 2.3.4.4b H5N1 viruses remain a top priority to mitigate their potential impact on public health and the agricultural sector.

## Supporting information

Graphical Abstract

## Funding

The work was funded by the UK Research and Innovation (UKRI), Biotechnology and Biological Sciences Research Council (BBSRC) and Department for Environment, Food and Rural Affairs (Defra, UK) research initiative ‘FluMAP’ and FluTrailMap-Avian grants (BB/X006166/1, BB/Y007298/1, BB/X006204/1 BB/Y007271/1)], the Pirbright Institute strategic program grant (BBS/E/PI/230001B, BBS/E/PI/230001C, BBS/E/PI/230002B, BBS/E/PI/230002C), BBS/E/PI/23NB0004, BBS/E/PI/23NB0003], the Medical Research Council (MRC) and Defra research initiative ‘FluTrailMap-One Health’ grant (MR/Y03368X/1) and The Global Challenges Research Fund (GCRF) One Health Poultry Hub grant (BB/S011269/1). The funders had no role in study design, data collection, data interpretation or the decision to submit the work for publication.

### Acknowledgement

We gratefully acknowledge U.S. Department of Agriculture (USDA), U.S. Centers for Disease Control and Prevention (CDC), the World Health Organization (WHO), Iowa State University, and St. Jude Children’s Research Hospital for sharing cattle virus sequences on Sequence Read Archive (SRA) on National Center for Biotechnology Information (NCBI). We gratefully acknowledge all data contributors, i.e., the authors and their originating laboratories responsible for obtaining the specimens, and their submitting laboratories for generating the genetic sequence and metadata and sharing via the GISAID Initiative, on which this research is based. All submitters of the data may be contacted directly via the GISAID website (https://www.gisaid.org). We thank Dr. Steve Martin for sharing Octet analysis software for receptor binding analysis. The graphical abstract was created using Biorender (https://www.biorender.com).

---

**Table S1.**
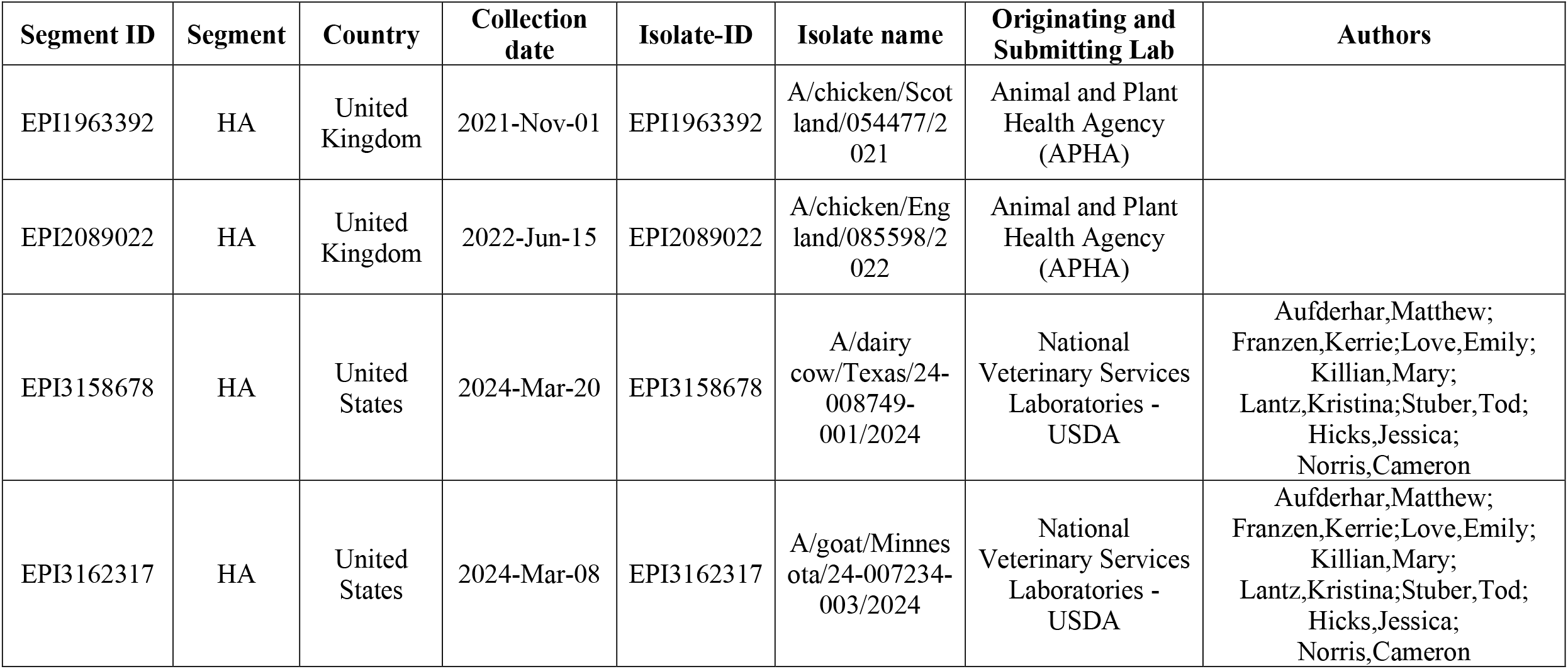
Authors, originating and submitting laboratories of the sequences from GISAID used in this study.

**Figure S1.**
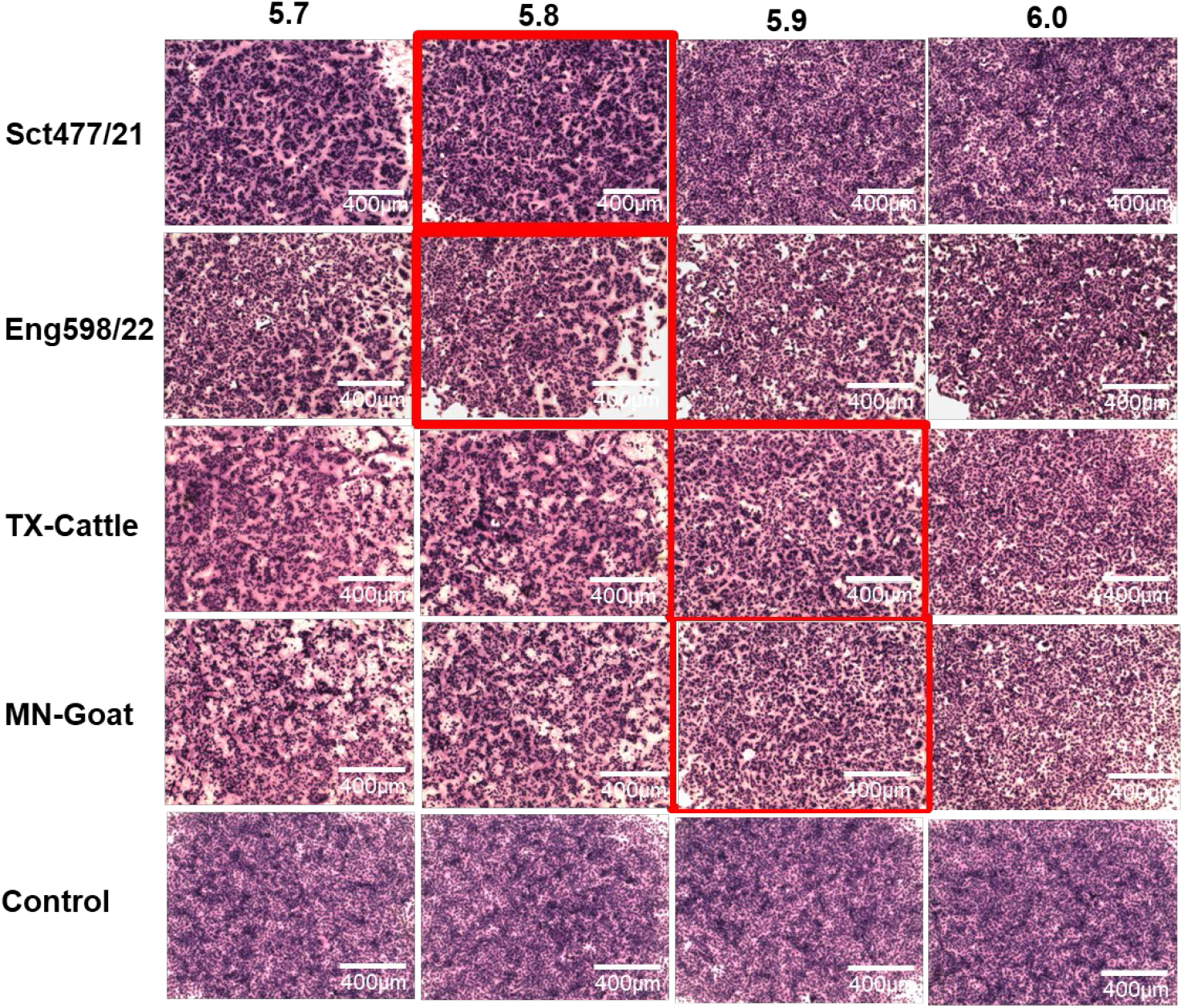
pH fusion of the panel H5N1 viruses. Monolayered Vero cells were infected with the panel viruses and subsequently treated with PBS at the indicated pH. Cells were fixed in acetone:methanol (1:1) and stained with Giemsa solution. Images were captured at a scale of 400 μm.

**Figure S2.**
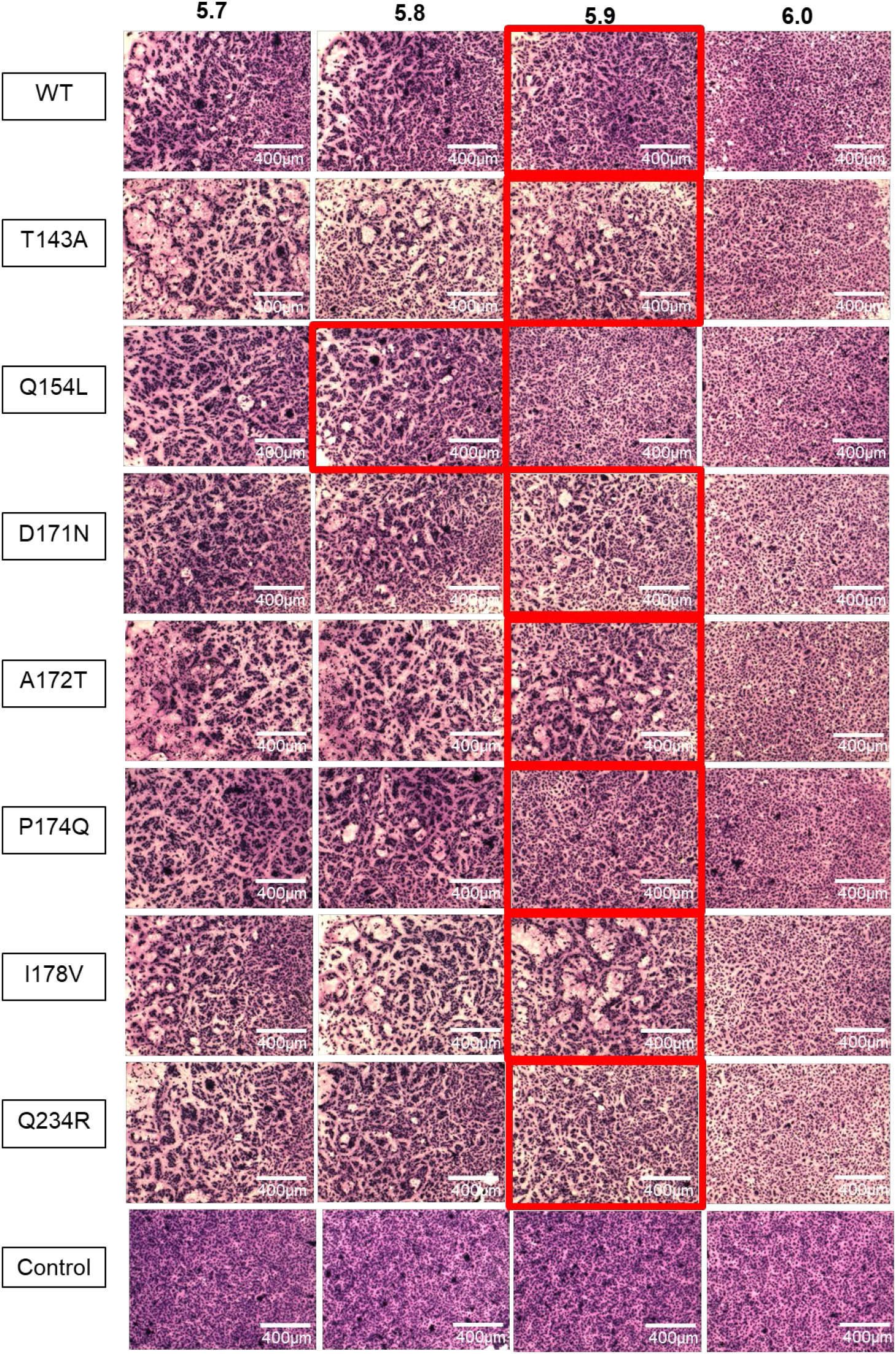
pH fusion of the H5N1 mutant viruses. Monolayered Vero cells were infected with the mutant viruses and subsequently treated with PBS at the indicated pH. Cells were fixed in acetone:methanol (1:1) and stained with Giemsa solution. Images were captured at a scale of 400 μm.

