## Supplementary figures and images for "The Haemagglutinin Gene of Bovine Origin H5N1 Influenza Viruses Currently Retains an Avian Influenza Virus phenotype"

### Graphical Abstract

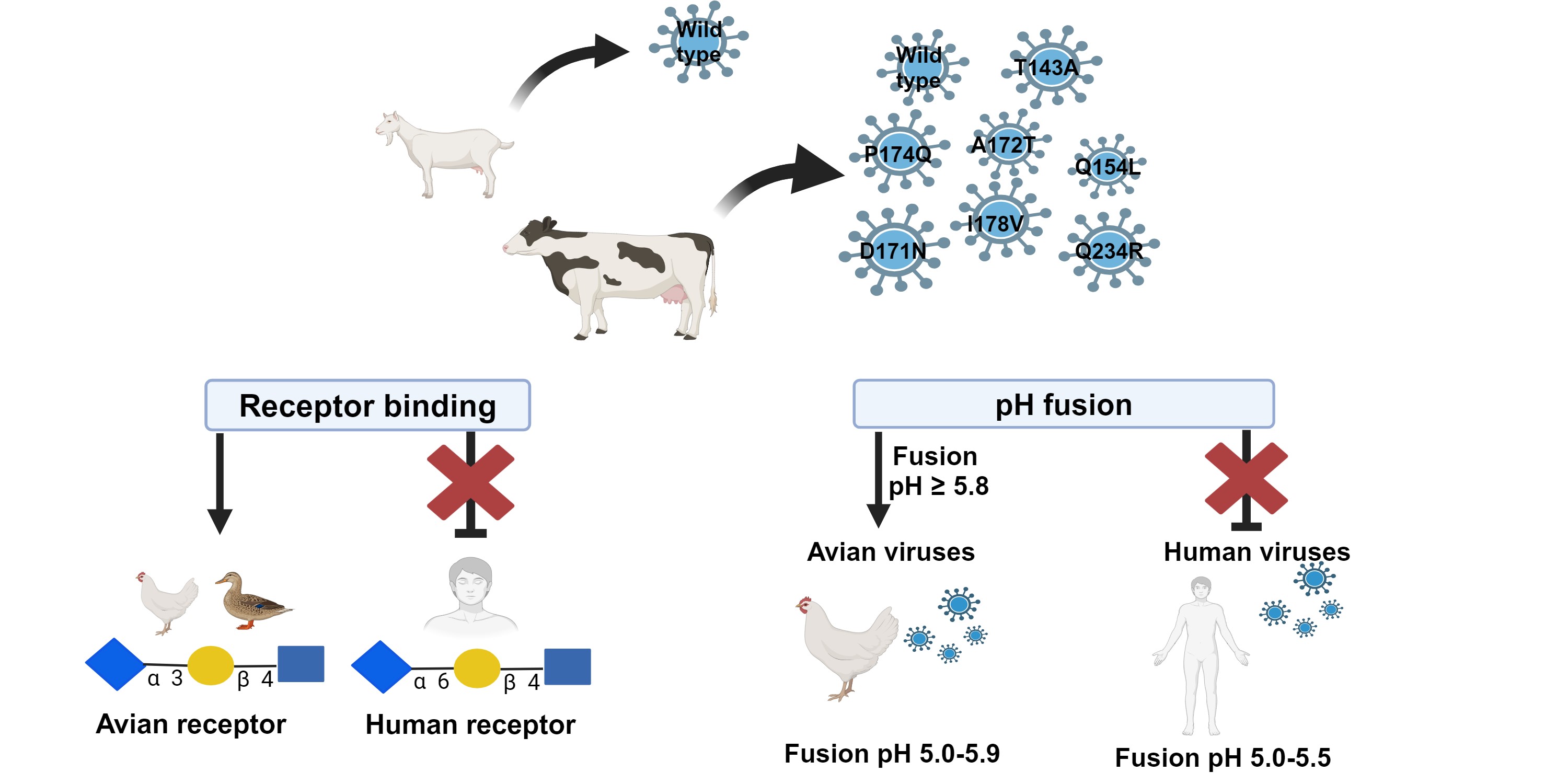
